# Comparison of Transcranial Doppler Ultrasound with Computational Fluid Dynamics: Responses to Physiological Stimuli

**DOI:** 10.1101/2022.02.22.480470

**Authors:** Harrison T. Caddy, Hannah J. Thomas, Lachlan J. Kelsey, Kurt J. Smith, Barry J. Doyle, Daniel J. Green

## Abstract

Few studies have compared transcranial Doppler (TCD) ultrasound with independent techniques such as computational fluid dynamics (CFD) simulations, particularly in response to stimuli. We compared TCD cerebral blood flow velocity in healthy participants with subject-specific CFD simulations to determine differences in techniques. Twelve participants underwent head and neck imaging with 3 Tesla magnetic resonance angiography. Velocity waveforms in the middle cerebral artery (MCA) were measured with TCD while velocity and diameter in the neck arteries were measured with duplex ultrasound at rest, hypercapnia and exercise. Subject-specific CFD simulations were developed for each condition, with velocity waveforms extracted in the same region as TCD. We found that absolute TCD velocities were significantly higher than CFD data, and non-significantly correlated across all conditions (r range 0.030-0.377, all P>0.05). However, relative changes from rest to hypercapnia and exercise generally exhibited significant positive correlations (r range 0.448-0.770), with the strongest correlation being average velocity change from rest to exercise (r=0.770, P<0.01). We have found that although absolute MCA velocity measurements from different sources vary, relative velocity changes yield stronger correlations regardless of source. Our findings indicate relative responses to physiological stimuli, along with absolute data, should be considered for analyzing cerebral blood flow velocity.

## INTRODUCTION

Transcranial Doppler (TCD) is an ultrasound technique which is commonly used for measuring blood flow velocities within the brain. It is typically used to insonate the middle (MCA), posterior (PCA) or anterior cerebral arteries via either the transtemporal, transorbital or suboccipital windows. A conventional TCD system consists of a low frequency 2 MHz probe and is either held in location by an operator, or secured using a head brace ^42^. TCD is beneficial for measuring cerebral blood flow velocity because it is non-invasive, reproducible, can be quickly performed in real-time, is portable and possesses high temporal resolution ^42^. Despite these benefits, results are known to be operator dependent ^25^, as well as limited by fixed transorbital windows and the probe frequency required to transit the bony skull. Furthermore, TCD can only measure velocity, as B-mode images of diameter cannot be simultaneously measured due to resolution constraints ^42^.

TCD has been used to measure blood flow velocity and derived metrics of arterial pulsatility and resistance in patients with cerebrovascular diseases ^4, 13, 18^, as well as being used in estimating cerebral blood flow (CBF) to different regions of the brain ^24^. It has also been used to assess prognosis and diagnosis of life-threating conditions such as sickle cell anemia, stenosis, hemorrhage and ischemia across a spectrum of patient ages ^25, 29^. TCD is also used extensively in healthy populations to better understand cerebrovascular responses to physiological stimuli including hypercapnia, hypocapnia, exercise, vessel occlusion, shear-mediated endothelial responses and neurovascular coupling ^5, 11, 15, 28, 41^. In addition, measurement of blood flow velocity using TCD has been used to infer volumetric cerebral blood flow – however the absence of diameter measurement is a limitation in this regard. Comparisons between volumetric flow measures in the internal carotid artery (ICA) and MCA velocities during the vasodilatory stimuli of hypercapnia ^43^ and exercise ^39, 40^, suggest intracranial diameter changes may result in underestimation of CBF when estimated from TCD ultrasound. There is also some evidence which suggests that cerebral artery diameter may vary within the cardiac cycle ^1, 7^ and that these arteries are also known to vasoconstrict in response to changes in arterial blood pressure in the context of cerebral autoregulation ^3, 11^. Consequently, using TCD velocities as a measure proportional to cerebral blood flow remains controversial in the absence of a B-mode image of diameter change ^1^. In some instances, this limitation has been addressed by using ICA or vertebral arteries (VA) as surrogates for intra-cranial vessels, which are contiguous with arteries such as the MCA and PCA, and can be imaged using real time duplex ultrasound.

Few studies have compared measurements of blood flow velocity obtained using TCD with alternative and independently derived techniques. One such alternative method for measuring blood flow within the brain is through magnetic resonance imaging (MRI), such as magnetic resonance angiography (MRA). Despite limitations in temporal resolution, this approach possesses high spatial resolution and is able to capture the complex three dimensional (3D) nature of cerebral arteries ^1^. Computational fluid dynamics (CFD) simulations provide a method for combining the strengths of both MRA and duplex ultrasound to calculate fluctuations of blood flow velocity within the cerebral vasculature with high spatial and temporal resolution. Although CFD has been used in conjunction with TCD methods for independent comparison ^14, 27, 30, 35^ under static conditions, few studies have investigated velocities measured using TCD with independent methods, such as CFD, in response to common physiological stimuli ^11^.

In this study, we tested the change in TCD velocity measurements in response to physiologically relevant stimuli such as hypercapnia and exercise, and compared these data to changes in simulated cerebrovascular velocities calculated using CFD methods. Our CFD approach combined ICA and VA duplex ultrasound measurements to prescribe volumetric flow inlet boundary conditions, and used 3D geometry based on individualized MRA-derived cerebrovascular reconstructions. We hypothesized that TCD- and CFD-derived velocity metrics would be similar and highly correlated at rest and in response to physiological stimuli.

## MATERIALS AND METHODS

### Participant Cohort and Medical Imaging

The experimental procedures used in this study were approved by The University of Western Australia Human Research Ethics Committee. A total of twelve healthy participants (6 female, 6 male) were recruited for this study with ages ranging between 19 and 28 years old. Participants were made aware of the experimental procedure and associated risk. Written consent was obtained for each participant prior to commencement of the experimental study. Prior to cerebrovascular stimuli, each participant underwent a 3 Tesla time-of-flight (3T TOF) MRA (Siemens MAGNETOM Skyra) neck and head scan. This scan had a pixel size of 0.31 mm and a slice thickness of 0.75 mm.

### Cerebrovascular Stimuli Procedure

Participants were exposed to conditions previously described by Thomas *et al*. ^41^, consisting of rest (5 minutes recumbent), hypercapnia (5 min of 6% CO_2_ via Douglas bag recumbent) and submaximal exercise (5 min recumbent cycling at 90 Watts) conditions respectively. Each session was conducted in the morning for all participants, who were instructed to fast (no food, tea or coffee) and abstain from consumption of alcohol or performing exercise in the 24 hours prior to the session. After each of the exposure condition durations, we simultaneously measured velocity (Doppler ultrasound) and intraluminal diameter (B-mode ultrasound) in the left and right ICAs and VAs using two identical 10-MHz linear array probes and high-resolution ultrasound machines (Terason 3200, Teratech, Burlington, MA) using standardized search techniques ^42^. Continuously during each condition, we measured the peak velocity envelope in the MCA, specifically in the right M1 segment via the middle transtemporal window using a 2-MHz TCD probe (TCD, Spencer Technologies, Seattle, WA) which was held in place with a headpiece (M600 bilateral head frame, Spencer Technologies) as per methods described previously ^15^. Throughout all exposure conditions, we continuously measured end-tidal partial pressure of CO_2_ and O_2_ using a gas analyzer (Gas Analyzer, ADInstruments, New South Wales, Australia). For analysis and calculation of ICA and VA flow and TCD MCA velocities, we considered data sampled within the final 30 second period of each 5 minute exposure condition. To mitigate cerebral priming, participants underwent a 10 minute washout period, remaining in a recumbent position, between each exposure condition, which was sufficient to return metrics (mean arterial pressure, heart rate, end tidal CO_2_, average MCA velocity from TCD) to baseline levels across all participants.

### MRA Reconstruction and Ultrasound Analysis

#### MRA 3D Reconstruction

To create the 3D fluid domain for the CFD simulations, we imported the DICOM images from the MRA scan into in-house image reconstruction software. We used region-growing techniques to select similar intensity labeled pixels greater than an intensity value of 200 to outline the fluid contained within the cerebrovascular geometry. The software uses an inbuilt marching cubes algorithm to create a 3D isosurface. This isosurface was globally smoothed to within 5% of the starting volume, which was then imported into STAR-CCM+ (v12, Siemens, Munich, Germany) to perform surface repair, remove reconstruction artifacts and perform local smoothing. Outlets were truncated perpendicular to the vessel centerline at least two bifurcations downstream from the Circle of Willis (CoW).

#### Duplex and TCD Ultrasound

The duplex ultrasound measurements of diameter and velocity at the ICAs and VAs over three cardiac cycles ^20^ were converted to time varying volume flow rate and waveform averaged and processed as described previously ^41^. This resulted in spline fitted and peak aligned data. Similarly, TCD maximal velocity waveform data from three cardiac cycles within the right M1 segment (Figure 1) were combined into an average waveform using the same process for each participant.

**Figure 1.**
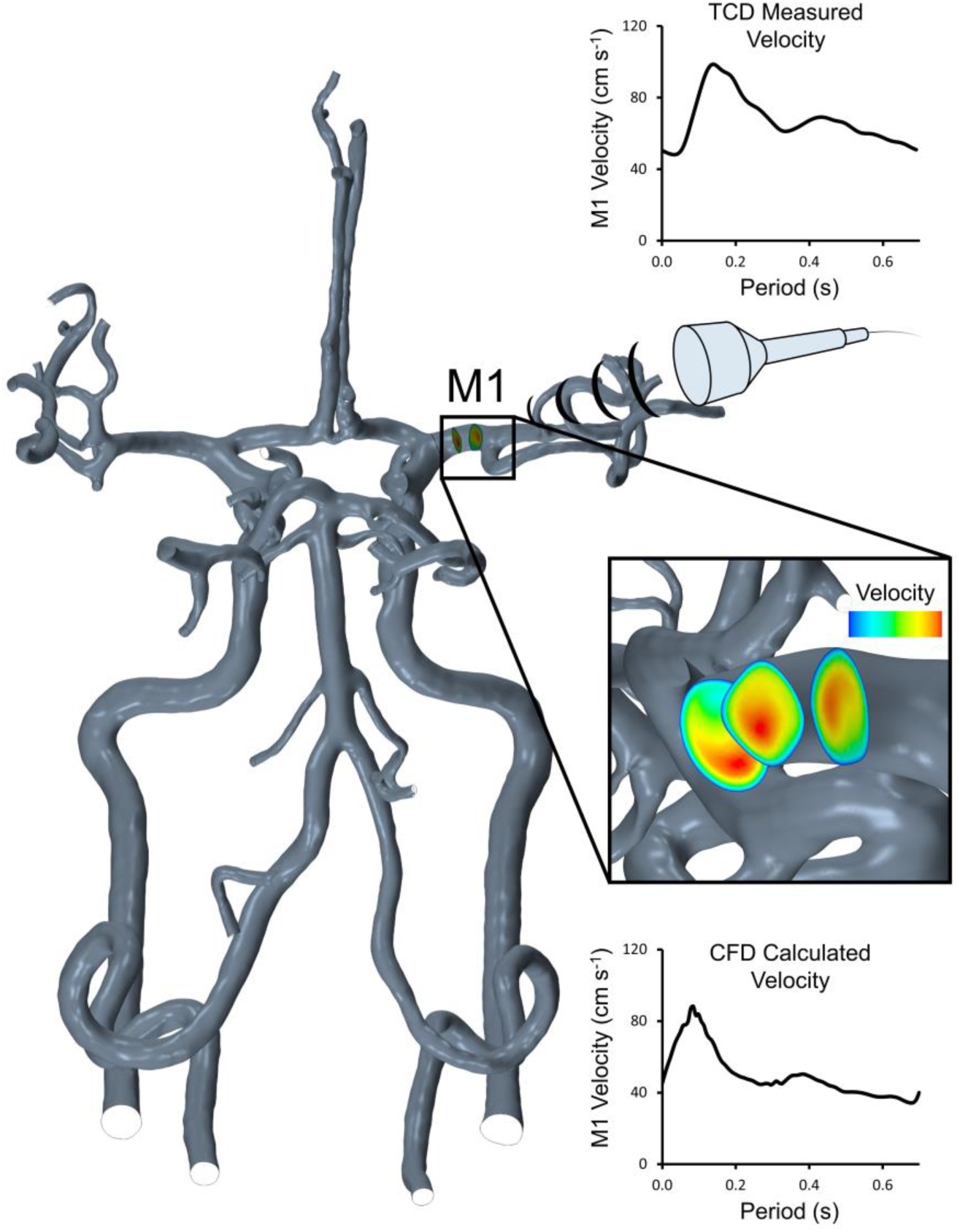
Example TCD probe (top) and consecutively spaced CFD constrained planes (bottom) demonstrating sampling of data within the right M1 segment. Within the CFD simulations, maximal velocity was calculated and extracted at each of these planes and averaged into a mean maximal value representative of the peak velocity envelope from TCD insonation of the right M1 segment in each participant. Example velocity waveforms from TCD and CFD sources from an individual are provided. Characteristics from these velocity waveforms were then extracted and compared between sources across a range of different exposure conditions (rest, hypercapnia and exercise).

### Computational Fluid Dynamics

#### Computational Mesh

The simulations were developed in the commercial CFD package STAR-CCM+. We used a combination of a polyhedral element mesh for the core of the fluid domain and 20 prism layer elements in the near wall boundary to sufficiently capture the velocity gradients and ensure accurate calculation of wall shear stress. In addition, we prescribed extrusions at the fluid boundaries equal to 11 times the boundary diameter to ensure adequate development of parabolic flow upstream and downstream of the fluid domain ^6^. Mesh core density was set proportional to local vessel diameter and the mesh settings were prescribed as per previously published work ^41^. These settings had been optimized to ensure mesh independence using the grid convergence index ^31^ and used the subject with the greatest inlet velocity measurements for this optimization. Optimal mesh size was deemed sufficient when the grid convergence index for wall shear stress within the CoW was found to fall below 3% ^41^. Final mesh sizes ranged from 9.1 to 16.7 million cells per geometry.

#### Boundary Conditions

Boundary conditions followed methods as described previously ^41^. Briefly, the three cardiac cycle duplex ultrasound measurements of velocity and diameter obtained at the ICAs and VAs were converted from the calculated volumetric flowrates to mass flow waveforms, assuming a fluid density of 1050 kg m^-3 22^. The measured velocity waveforms for the rest, hypercapnia and submaximal exercise conditions were all processed using this method and prescribed at the corresponding extruded inlet using a plug flow condition, which developed into a parabolic profile throughout the extruded inlet region before entering the subject specific cerebrovascular geometry. Outlet boundary conditions were implemented using the same WALNUT code described previously ^41^, which initially splits blood flow exiting the fluid domain into seven regions (left and right posterior, left and right middle, anterior, cerebellum and ophthalmic arteries) and accounts for the presence or absence of communicating arteries in the CoW, with different flow distributions to these regions based on the average volumetric flow measured from the ICAs and VAs. Within these regions, flow was then split using an adaptation of the Murray’s law formulation ^10^ as defined in equation 1, using an exponent of n=2.33 ^41^.

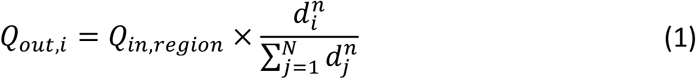

Where *Q*_*out,i*_ is the flow out of outlet *i, Q*_*in,region*_ is the WALNUT calculated flow into the region of the cerebrovasculature which is shared by outlet *i* as described previously ^41^, *d* is outlet diameter and *n* is the flow split exponent.

#### Physical Assumptions

Blood was assumed incompressible with a density of 1050 kg m^-3 22^. The non-Newtonian nature of blood viscosity was modelled using the Carreau-Yasuda viscosity model using parameters appropriate for blood flow within the CoW ^19^. We assumed the arterial walls were rigid, with a no-slip boundary condition and a laminar flow regime in line with previous cerebrovascular simulations ^2, 19, 36, 41^.

#### Simulation Execution

Simulations ran for three consecutive cardiac cycles to ensure flow stabilization, with results extracted from the fourth cardiac cycle. The implicit unsteady segregated flow solver was employed, which uses the Semi-Implicit Method for Pressure Linked Equations (SIMPLE) for coupling the velocity and pressure components of the Navier-Stokes equations. We used second-order temporal discretization with an automated time-step control which permitted the time-step to vary between 0.001 and 0.005 s depending on the Courant number. New time-steps were triggered if absolute continuity and momentum residuals fell below a value of 10^−9^, or if the number of inner iterations reached 50. Our simulations were executed using the STAR-CCM+ finite-volume method on Magnus, a Cray XC40 supercomputer (Pawsey Supercomputing Centre, Perth, Australia) housing a total of 1488 compute nodes each containing 24 cores per node. We ran each simulation utilizing 25 nodes over a collective of 500 cores. The rest, hypercapnia and exercise simulations required an average of 3000, 2800 and 2000 core hours respectively to run, which equated to total simulation run times ranging from approximately 3 to 7 h.

### Data Collection, Analysis and Statistics

Simulations were allowed to stabilize over three cardiac cycles, after which we extracted maximal velocity waveforms from each simulation over the fourth cardiac cycle using the average of the maximum value from three consecutively spaced constrained planes located within the right M1 segment (Figure 1) to represent the region insonated using TCD. The velocity waveforms extracted from the simulation and from the TCD ultrasound envelope for each participant and exposure condition were analyzed for their characteristics using a custom MATLAB script (R2016, Mathworks, Natick, MA). We extracted the systolic, average and end-diastolic maximal velocities from these waveforms. We used paired t-tests for comparison of distributions and Pearson’s correlation and Bland-Altman plots to investigate the correlations between absolute and relative change in CFD and TCD data for each participant in response to different stimuli. Normality was tested using the Shapiro-Wilk test. Where applicable, data is presented as group mean ± standard deviation. Statistical significance was assumed for p-values where P<0.05.

## RESULTS

### Participant Cardiorespiratory Responses and MRA Reconstructions

Participants had an average age, BMI and VO_2_max score of 22.9±3.4 years, 21.6±2.9 kg m^-2^ and 44.7±9.4 mL kg^-1^ min^-1^ respectively. Participant cardiorespiratory responses including end tidal CO_2_, blood pressure, heart rate as well as ultrasound derived average volumetric blood flow at the ICAs and VAs under resting, hypercapnia and exercise conditions are displayed in Table 1. We reconstructed 3D models of each of the twelve cases from the MRA data collected, which all exhibited a complete Circle of Willis (see Supplementary Material, Figure S1).

**Table 1.** Participant cardiorespiratory responses under different conditions.

| | Rest<br>(Average ± SD) | Hypercapnia<br>(Average ± SD) | Exercise<br>(Average ± SD) |
| --- | --- | --- | --- |
| End-tidal $\text{CO}_2$ (mmHg) | $42.2 \pm 2.5$ | $50.7 \pm 1.9$ | $43.8 \pm 4.2$ |
| Mean arterial pressure (mmHg) | $100.0 \pm 8.1$ | $106.0 \pm 11.8$ | $124.8 \pm 15.2$ |
| Heart rate (bpm) | $68.7 \pm 9.5$ | $74.1 \pm 8.1$ | $120.4 \pm 21.3$ |
| Mean blood flow ($\text{mL min}^{-1}$) | | | |
| LICA | $268.9 \pm 63.3$ | $374.6 \pm 79.3$ | $272.8 \pm 67.2$ |
| RICA | $218.8 \pm 74.2$ | $308.6 \pm 79.3$ | $262.9 \pm 50.3$ |
| LVA | $74.2 \pm 26.4$ | $119.4 \pm 30.9$ | $79.7 \pm 27.1$ |
| RVA | $56.2 \pm 18.3$ | $97.0 \pm 32.3$ | $68.4 \pm 34.0$ |

Data are mean ± standard deviation. LICA = left internal carotid artery; RICA = right internal carotid artery; LVA = left vertebral artery; RVA = right vertebral artery.

### Absolute Velocity Data Distributions at Single Conditions

In general, measurements of maximal systolic, average and end diastolic velocity were all significantly higher (all P<0.05) in the TCD data compared to the CFD simulations (Figure 2). The mean of the systolic velocity distribution ranged from 33-73% higher in the TCD data compared to CFD across the exposure conditions. Greater differences in means were observed for average velocity, with increases ranging from 62-85% when comparing TCD to CFD data across all exposure conditions. We observed the highest changes in end diastolic velocity, with increases ranging from 85-106% between TCD and CFD data across all conditions.

**Figure 2.**
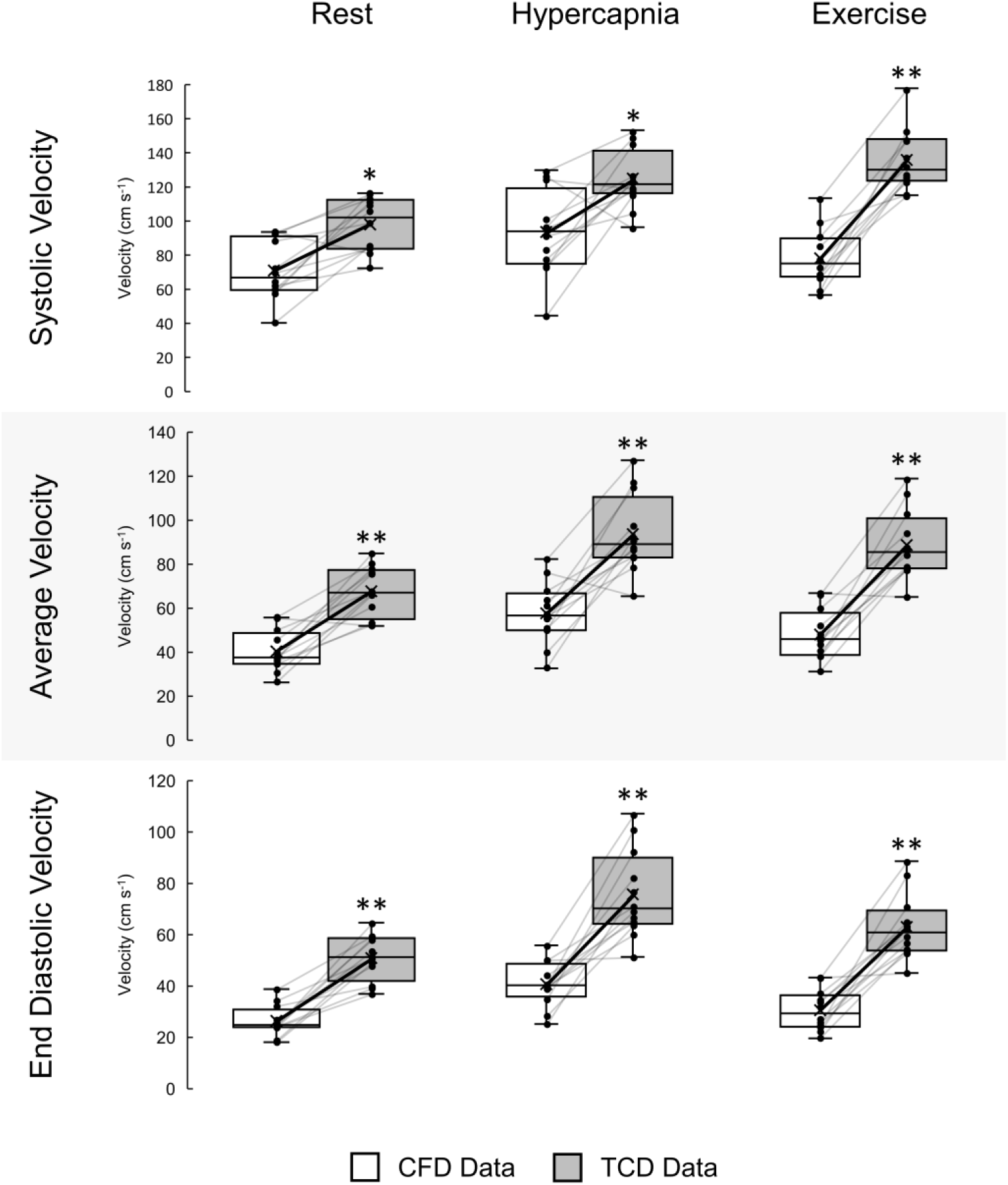
Box plot distributions of systolic, average and end diastolic maximal velocity extracted from CFD (white; n=12; 6 male, 6 female) and three cycle averaged TCD (grey; n=12; 6 male, 6 female) data in the right M1 segment. Individual differences between CFD and TCD data are presented as black dots with grey connecting lines. The solid black line connecting the cross (X) in each box indicates the changing means of the distributions. These data were collected for each of the stimuli conditions of rest, hypercapnia and exercise. Stars (*) indicate the level of significance (*P<0.05; **P<0.001) using t-tests between CFD and TCD data.

### Relative Velocity Data Distributions in Response to Physiological Stimuli

We investigated the relative change distributions of velocity waveform characteristics from rest to hypercapnia and rest to exercise from the CFD and TCD datasets (Figure 3). We found that the relative change in velocity metrics from rest to hypercapnia were similar between CFD and TCD (all P>0.05), with the means of CFD relative velocity changes found to be slightly higher than the changes from TCD data for systolic (34±30% vs 28±12%, P=0.579), average (44±26% vs 39±15%, P=0.532) and end diastolic velocity (58±33% vs 49±22%, P=0.477). A relative change in systolic velocity from a CFD case in response to hypercapnia was also observed to be greater than the higher quartile range limit. While small differences between CFD and TCD distributions were observed for average (22±27% vs 32±20%, P=0.323) and end diastolic (19±29% vs 26±24%, P=0.566) velocities, the change in systolic velocity from rest to exercise was significantly higher in the TCD data compared to CFD (40±18% vs 14±22%, P=0.006).

**Figure 3.**
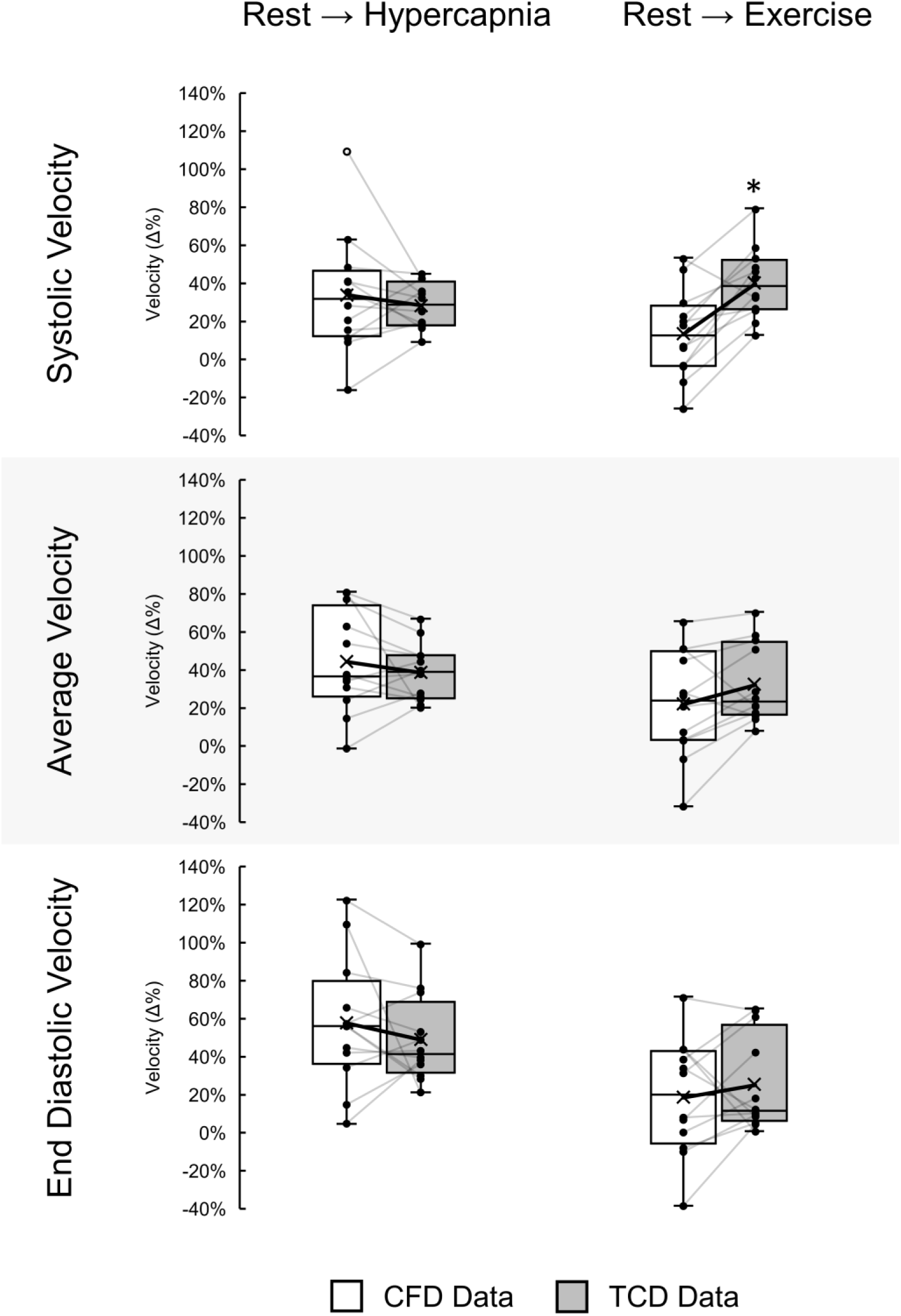
Box plot distributions of relative change (Δ%) in systolic, average and end diastolic maximal velocity extracted from CFD (white; n=12; 6 male, 6 female) and three cycle averaged TCD (grey; n=12; 6 male, 6 female) data between the conditions of rest to hypercapnia and rest to exercise in the right M1 segment. Individual differences between CFD and TCD relative change data are presented as black dots with grey connecting lines. The solid black line connecting the cross (X) in each box indicates the changing means of the distributions. A CFD data point of relative change in systolic velocity from rest to hypercapnia that is outside the higher quartile range limit is displayed as a hollow circle. Stars (*) indicate the level of significance (*P<0.05; **P<0.001) using t-tests between CFD and TCD data.

### Absolute Velocity Data Correlations at Rest and Physiological Stimuli

For systolic, average and end diastolic velocity under the conditions of rest (Figure 4), hypercapnia (Figure 5) and exercise (Figure 6), we found weak positive non-significant relationships (all P>0.05) that exhibited wide ranging limits of agreement and negative biases. Correlations between absolute velocity characteristics from CFD and TCD sources with total average flow from the ICAs and VAs across all stimuli were also computed (see Supplementary Material, Figure S2). Correlations of total average inlet blood flow to velocity from the CFD data were all significantly correlated (all P≤0.05), while TCD data yielded very mild non-significant correlations (all P>0.05), irrespective of exposure conditions or velocity metric.

**Figure 4.**
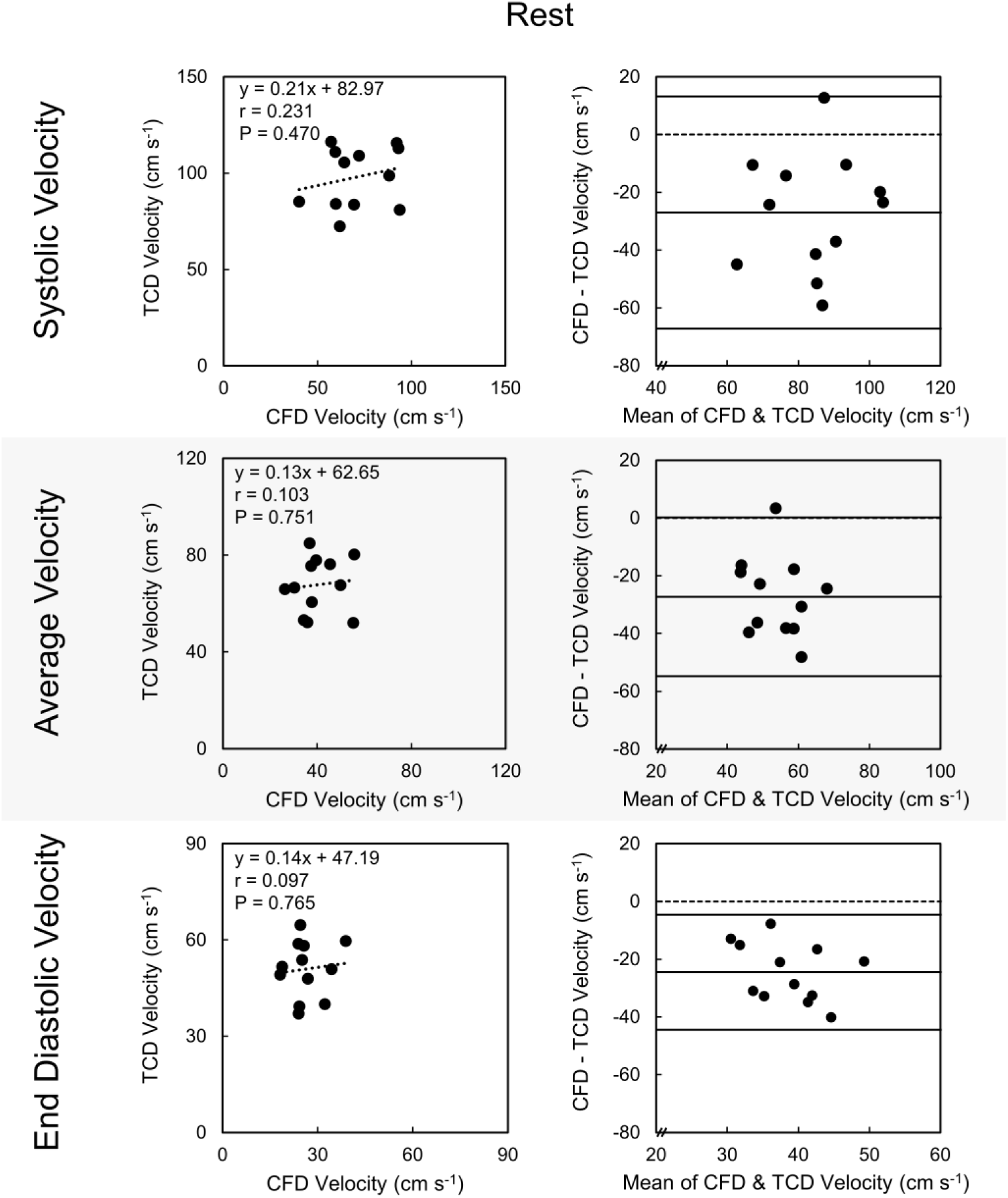
Correlation plots (left) and corresponding Bland-Altman plots (right) for absolute systolic, average and end diastolic maximal velocity extracted from CFD (bottom axis; n=12; 6 male, 6 female) and three cycle averaged TCD (left axis; n=12; 6 male, 6 female) data in the right M1 segment. These data are collected from the rest condition. The linear regression equation, Pearson’s correlation coefficient (r) and p-value (P) are displayed for each correlation plot.

**Figure 5.**
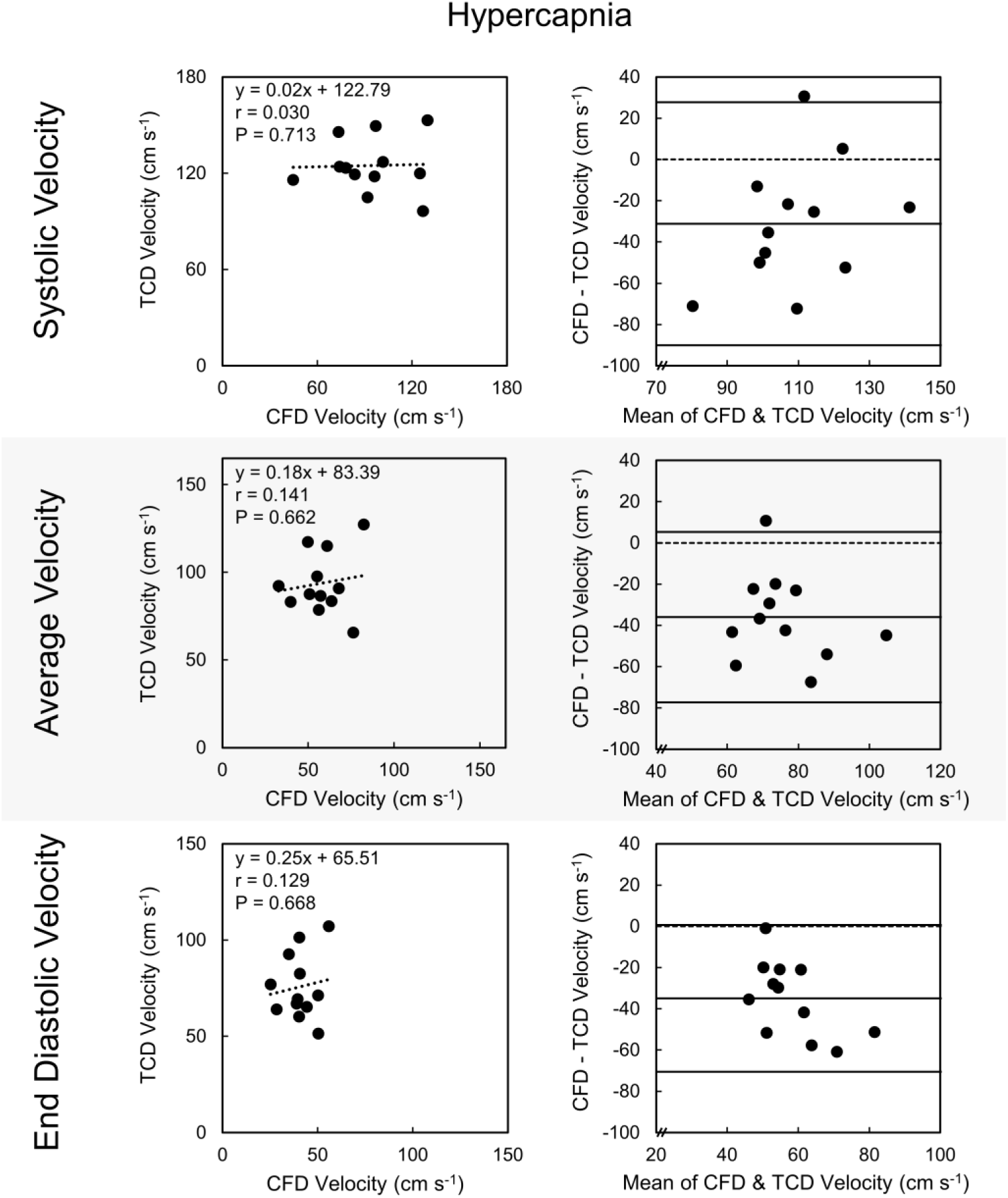
Correlation plots (left) and corresponding Bland-Altman plots (right) for absolute systolic, average and end diastolic maximal velocity extracted from CFD (bottom axis; n=12; 6 male, 6 female) and three cycle averaged TCD (left axis; n=12; 6 male, 6 female) data in the right M1 segment. These data are collected from the hypercapnia condition. The linear regression equation, Pearson’s correlation coefficient (r) and p-value (P) are displayed for each correlation plot.

**Figure 6.**
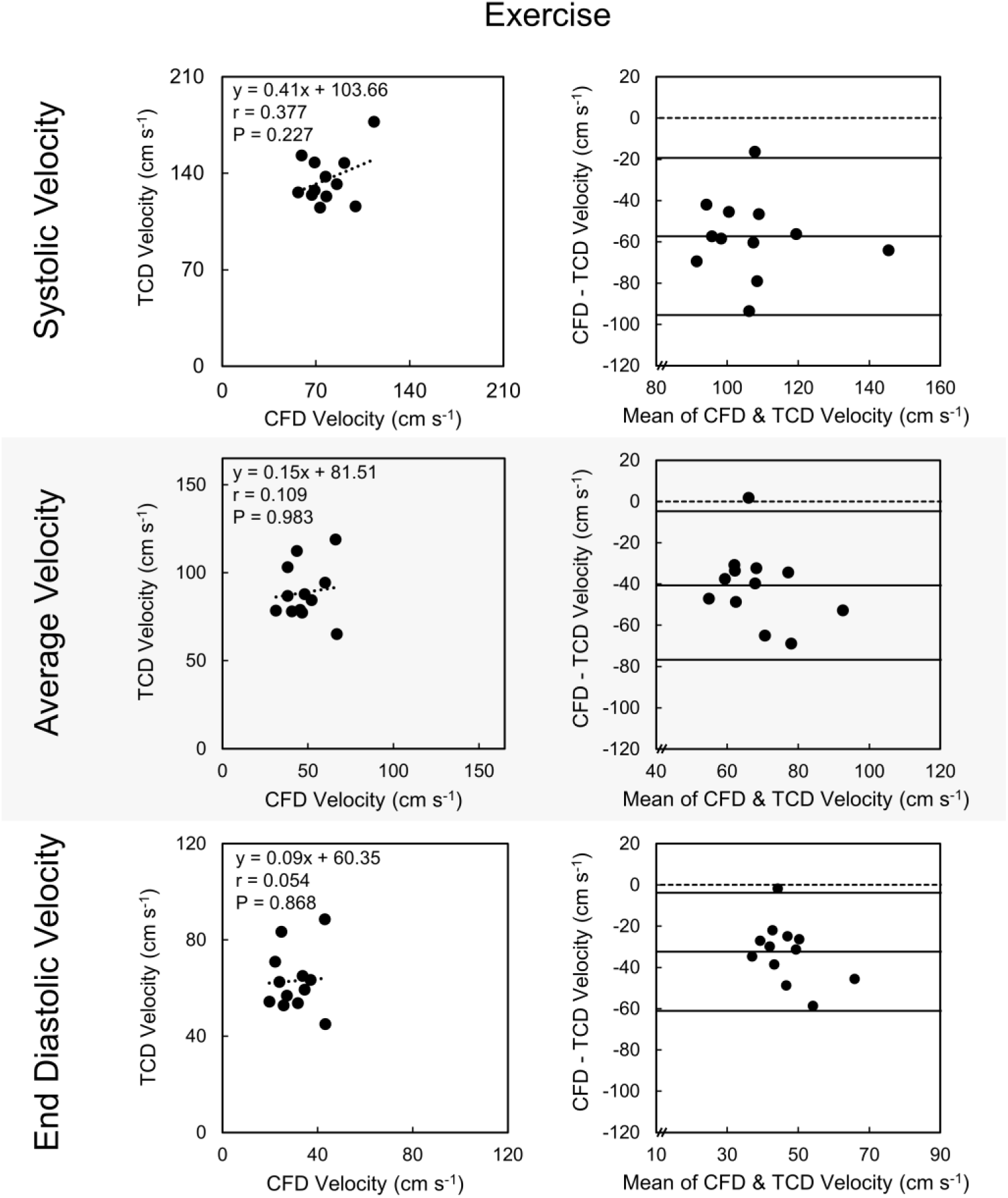
Correlation plots (left) and corresponding Bland-Altman plots (right) for absolute systolic, average and end diastolic maximal velocity extracted from CFD (bottom axis; n=12; 6 male, 6 female) and three cycle averaged TCD (left axis; n=12; 6 male, 6 female) data in the right M1 segment. These data are collected from the exercise condition. The linear regression equation, Pearson’s correlation coefficient (r) and p-value (P) are displayed for each correlation plot.

### Relative Velocity Data Correlations in Response to Physiological Stimuli

In comparison, correlations of CFD to TCD data for the relative change in velocity waveform characteristics of systolic, average and end diastolic velocity from rest to hypercapnia (Figure 7) and rest to exercise (Figure 8) yielded stronger moderate positive correlations, although limits of agreement remained large. Changes in systolic (r=0.588, P=0.04) and average (r=0.577, P=0.05) velocity from rest to hypercapnia were significantly correlated, with wide limits of agreement (−46% and 57%, -38% and 49%), while end diastolic velocity (r=0.448, P=0.14) was not significantly correlated and exhibited larger limits of agreement (−54% and 72%). Similarly, for the change from rest to exercise, relative changes in systolic (r=0.604, P=0.04) and average (r=0.770, P<0.01) velocity were found to be significantly correlated between CFD and TCD data, again with wide ranging limits of agreement (−64% and 11%, - 46% and 25%), while end diastolic (r=0.508, P=0.09) velocity was not significantly correlated and again displayed the largest range of limits of agreement (−62% and 48%). Excluding relative systolic velocity change bias from rest to exercise (−27%), the biases fell within ±10% for all other velocity metrics and exposure conditions. A mild positive proportional bias was observed across most relative velocity change metrics. We also investigated correlations between relative changes in velocity characteristics from CFD and TCD sources with relative change in total average flow from the ICAs and VAs in response to stimuli (see Supplementary Material, Figure S3). We again observed strong positive and significant correlations between average total inlet blood flow and the relative change in MCA velocity calculated from CFD simulations across both responses to hypercapnia and exercise (all P<0.05). Relative changes in TCD velocity metrics all remained positive, however, outside of average velocity from rest to exercise (r=0.588, P=0.044), these correlations remained mild and non-significant. Nonetheless, the mild positive correlations for relative change data were stronger across all relative change TCD velocity metrics and physiological stimuli compared to the correlations with inlet flow using absolute data.

**Figure 7.**
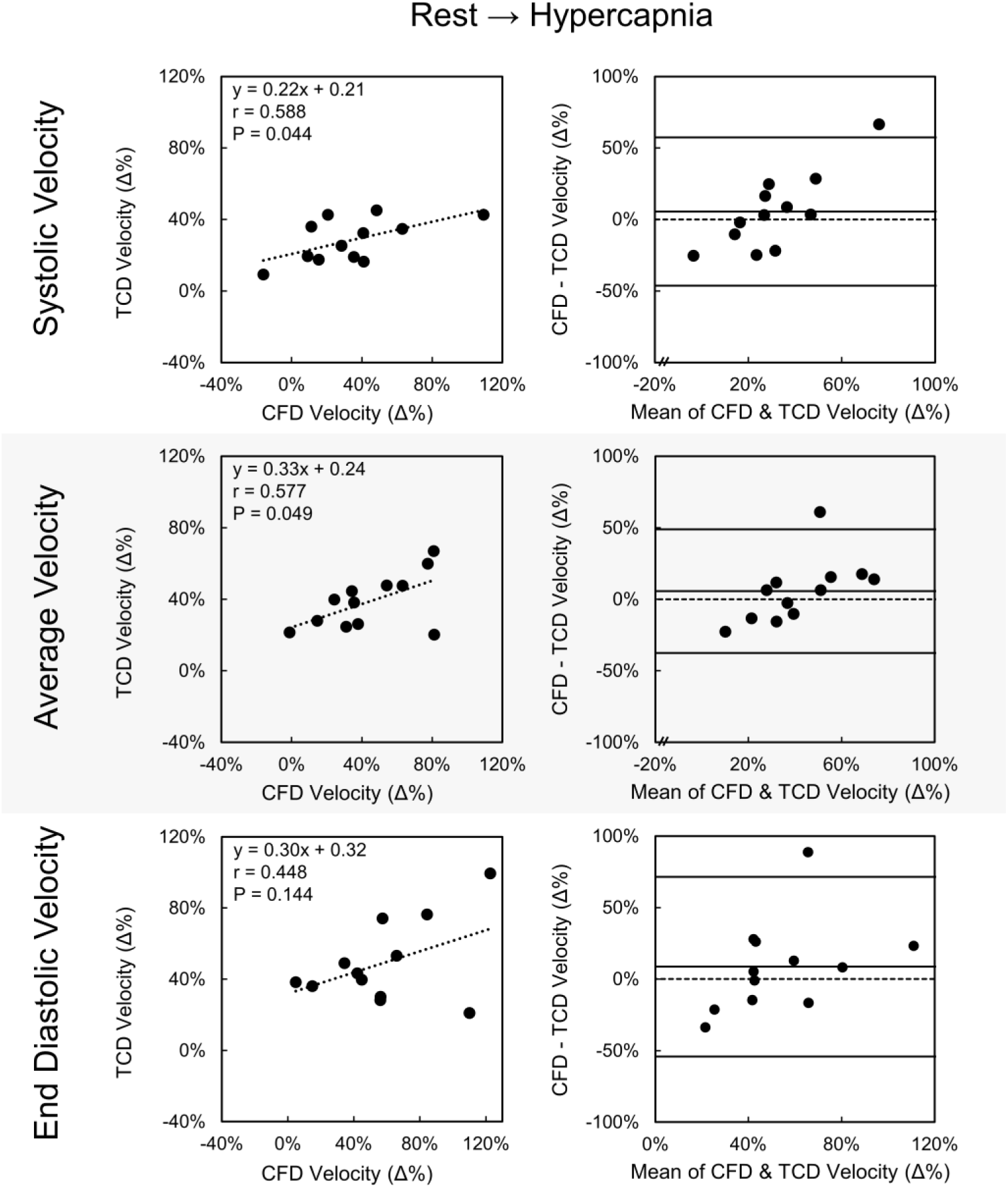
Correlation plots (left) and corresponding Bland-Altman plots (right) of the relative change (Δ%) in systolic, average and end diastolic maximal velocity extracted from CFD (bottom axis; n=12; 6 male, 6 female) and three cycle averaged TCD (left axis; n=12; 6 male, 6 female) data in the right M1 segment for rest to hypercapnia. The linear regression equation, Pearson’s correlation coefficient (r) and p-value (P) are displayed for each correlation plot.

**Figure 8.**
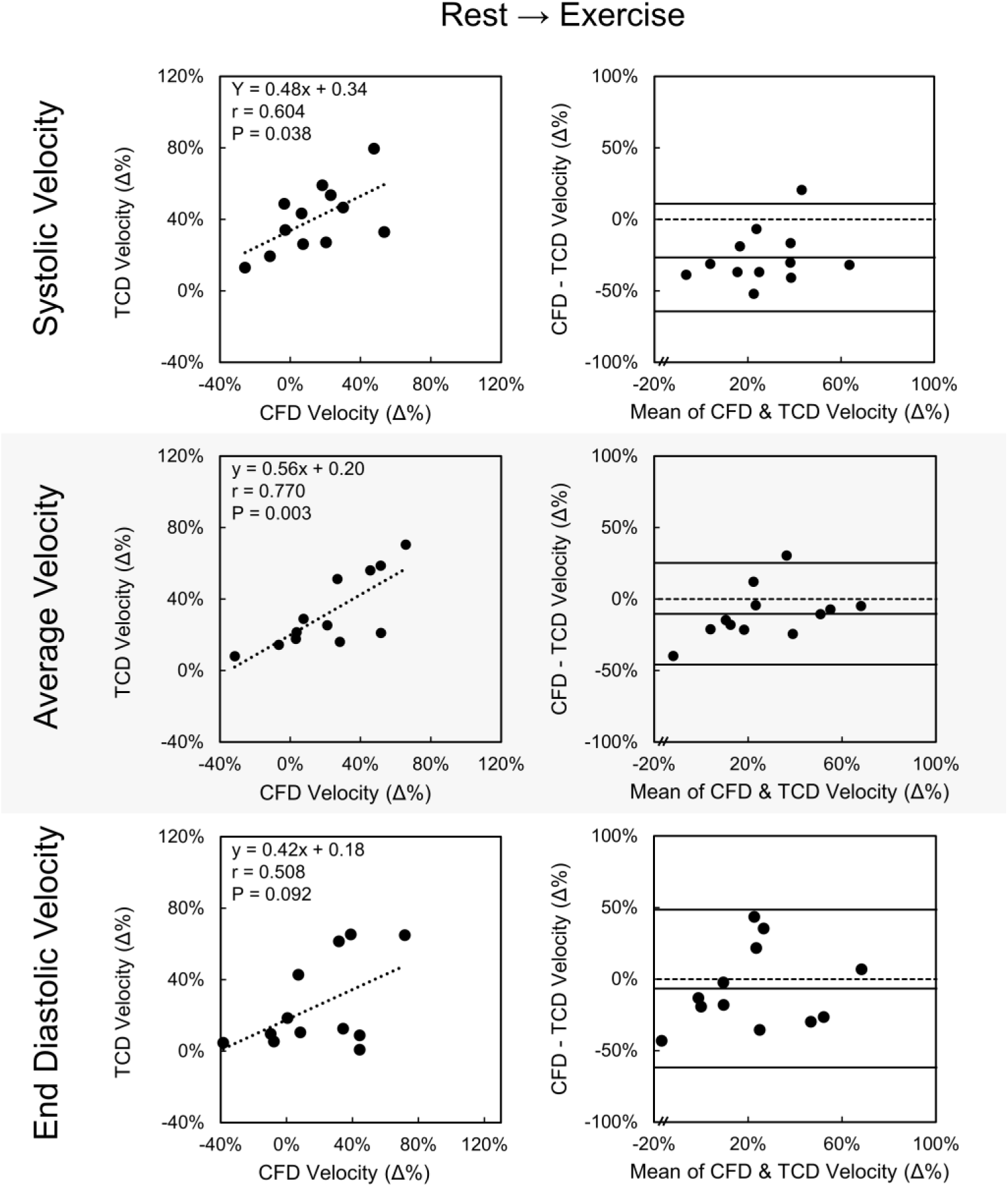
Correlation plots (left) and corresponding Bland-Altman plots (right) of the relative change (Δ%) in systolic, average and end diastolic maximal velocity extracted from CFD (bottom axis; n=12; 6 male, 6 female) and three cycle averaged TCD (left axis; n=12; 6 male, 6 female) data in the right M1 segment for rest to exercise. The linear regression equation, Pearson’s correlation coefficient (r) and p-value (P) are displayed for each correlation plot.

## DISCUSSION

In this study we investigated the differences in velocity obtained from two sources, subject-specific CFD simulations and TCD ultrasound, under the three exposure conditions of rest, hypercapnia and exercise. In contrast to our initial hypothesis, we found that absolute TCD velocities were significantly higher than those calculated using CFD, and that these comparative data sets were uncorrelated, irrespective of exposure condition. This disparity between velocity data could be explained by underestimation from CFD simulations, overestimation from TCD methods, or a combination thereof. Despite the differences in absolute data observed between TCD and CFD methods, changes in response to each physiological condition were more similar with moderate to strong correlation.

Comparison of relative changes in velocity may minimize the impact of systematic sources of variability, since relative changes could serve to reduce error through cancellation. In concert with this mathematical explanation, it is also plausible that physiological variability may be reduced when a standardized stimulus (hypercapnia, exercise) is applied, compared to the multiple physiological mechanisms that combine to influence absolute data. It is also important to note, however, that although bias was reduced and correlations improved in the response to stimuli, there was still some evidence of proportional bias in the Bland-Altman plots, along with variable limits of agreement. These data indicate there may be range of improved agreement between either sources for moderate individual increase responses to stimuli, as opposed to low or high responses. This is also reflected in greater agreement for the more moderate responses from rest to exercise, compared to the larger changes found with rest to hypercapnia. While both of these physiological stimuli are associated with some degree of hypercapnia, 6% CO_2_ exposure provides a much larger stimulus. Hypercapnia is a powerful cerebral vasodilator, inducing possible MCA diameter change. The assumption of fixed 3D geometry in CFD simulations may be associated with more error and variability in increased hypercapnic states, potentially explaining the higher correlations we observed in response to exercise than the 6% CO_2_ exposure condition. In any event, reporting relative changes in velocity in response to stimuli may be an important adjunct to comparison of absolute measures in future studies.

One explanation for the disparity between CFD and TCD estimates of absolute velocity is that CFD calculated velocities could be influenced by variability in the ultrasound derived input flows, collected in the ICAs and VAs. Although flow data derived from duplex ultrasound have been found to correlate with MR based estimates ^26^, variability in this data can be high due difficulties associated with neck insonation and operator-related issues. Interestingly however, ultrasound measurements have been found to *overestimate* blood flow velocity compared to MR imaging (30), indicating that low CFD data in the current study may not be ascribed to low ultrasound derived flow inputs alone. Furthermore, average ICA and VA ultrasound flows in our study were similar to those observed in previous experiments ^26, 32, 33, 37^. Whilst these findings suggest that the inputs to CFD were likely to be reliable, the combination of multiple data sources (ultrasound diameter and velocity, MRI-derived 3D geometry) along with CFD modelling assumptions may nonetheless have contributed to an underestimation of velocities using the CFD approach. Alternatively, an overestimation of velocity derived from the TCD approach may have occurred, due to phenomenon such as spectral broadening which can exaggerate peak blood flow velocity by up to 35% when lower megahertz probes are used ^12, 16^. The TCD and CFD data may also have differed due to the rigid wall modelling assumption used in CFD calculation, although the degree to which changes in MCA diameter impact velocity remains a matter of debate. However, as an example, the incorporation of a dilation of the MCA for a given inlet flow would likely further reduce the velocity calculated from the CFD simulations and exacerbate the discrepancies found in this study. Finally, we investigated the association between variables using correlation between inlet flow to the brain (derived from duplex ultrasound) and the absolute (see Supplementary Material, Figure S2) and relative change (see Supplementary Material, Figure S3) in MCA velocity calculated using either CFD simulations or directly measured with TCD. We found that CFD calculated absolute and relative change in velocity were significantly correlated with the inlet flows, whereas TCD velocity was mostly uncorrelated with duplex ultrasound inlet data, irrespective of the exposure conditions. Consequently, the CFD simulations are likely to reflect, but also potentially exacerbate any changes in the calculated inlet flows from the neck arteries. These data provide insight regarding the capacity of CFD and TCD to accurately reflect absolute brain blood flow in humans.

Although predicated on different hypotheses, two previous papers have compared TCD data with CFD simulations. Jahed *et al*. compared velocity measurements from two cerebral aneurysmal cases with corresponding TCD data in the anterior, middle and posterior cerebral arteries ^17^. Differences in velocity measurements were observed in the MCA between CFD and TCD sources and the authors concluded that TCD tests may introduce error and possibly lead to incorrect decisions regarding clinical diagnosis and treatment. CFD methods have also been used in the context of investigating sickle cell anemia ^30^. The findings indicated that misplacement of the TCD sampling and averaging region within a localized region of low or high velocity in the MCA could also lead to misdiagnosis. Our findings in 12 healthy subjects add to these previous experiments, in showing that TCD and CFD approaches to velocity assessment can differ, at least when absolute comparisons between derived velocities are compared.

Although validation of either dataset is difficult to ascertain, the question of whether TCD or CFD approaches provide a more accurate reflection of the true changes in velocity from physiological stimulation can be somewhat informed by consideration of previous studies which have compared techniques to magnetic resonance approaches. Seitz *et al*. found that TCD velocities exceeded MRA measured velocities by around 30% and reported low correlations between these approaches ^34^. Chang *et al*. also reported 30% greater velocities via TCD ^8^, but found that phase contrast MRA techniques correlated strongly with TCD. A study by Meckel *et al*. compared 4D phase contrast MRI (4D PCMRI) and transcranial color-coded duplex sonography ^23^ and also found TCD derived data was higher than 4D PCMRI, along with weak to mild correlations between these approaches. Leung *et al*. reported higher peak velocities using TCD than PCMRA ^21^ but also reported strong correlations when resting and hypercapnia-derived data were compared between approaches. Taken together, these studies indicate that, at rest and in response to physiological stimuli, TCD approaches may present higher velocities when compared to MR based methods, but the degree to which they correlate is variable. However, despite the variabilities between in sources, our results suggest that either technique (TCD or CFD) remain practical and beneficial in understanding the relative change in velocities in response to physiological stimuli in the brain.

Although we believe the methods used in this study to investigate comparisons between CFD and TCD data are rigorous, and note that they have been previously published ^41^, the present study is not without its limitations. Our cohort was limited to young, healthy individuals with no pre-existing cardiovascular diseases. Consequently, our findings may vary from some aforementioned studies due to their use of aging cohorts, or individuals with existing cardiovascular diseases or risk factors. The MRA scans that were segmented to produce the 3D models used in our study were performed with resting supine participants. Ideally, provided availability of specialized equipment, MRA scans should be captured in response to exposure conditions, allowing any changes in vessel diameter to be embedded in future CFD simulations. Ultrasound imaging was performed in a recumbent position during exposure conditions while a resting supine body position was required for the MRA scans. Although matching of imaging positions is preferable, performing ultrasound imaging outside of a recumbent position is difficult during exercise ^38^. Although care was made to ensure placement of sampling regions was consistent between TCD measurement and CFD simulations, this sampling was still operator dependent. While we used previously established methods ^20, 41^ for ultrasound waveform averaging, additional averaging of cardiac cycles may also serve to reduce the variability in results. In the CFD simulations, outlet boundary conditions were distributed using resting regional flow measurements derived from literature and diameter-based flow splitting exponents which were constant across all outlets. In the absence of regional brain blood flow data, particularly in response to stimuli, an exponent value appropriate for cerebrovascular vessels was used ^41^, however research has suggested that this exponent may vary for each individual outlet ^9, 10^. A localized outlet splitting method as described by Chnafa *et al*. ^10^, provided access to flow data in the brain, may be more appropriate in future CFD based cerebrovascular research. Our simulations employed rigid wall modelling as the resolution of the MRA data collected was unable to resolve arterial wall thickness and subject specific material properties were not known. Alternatively, implementation of fluid structure interaction (FSI) modelling would allow the vessel wall to deform and absorb energy throughout the cardiac cycle. However, this would likely reduce the velocities calculated from CFD simulation - further exacerbating the discrepancies observed between TCD and CFD velocities. True validation of CFD methods was unable to be performed in this study due to imaging limitations. Independently captured time varying image datasets using 4D MRI methods may help provide validation of future cerebrovascular CFD simulations. Finally, with only 12 participants, the number of cases investigated in this study is relatively small. However, despite the low number of cases, we still observed statistically significant results which may have important implications for future physiological research.

In conclusion, we aimed to compare velocity measurements obtained within the MCA under resting and external physiological stimuli (hypercapnia, exercise) conditions using TCD ultrasound and independently constructed CFD simulations. Although we found discrepancies between absolute velocity data obtained between CFD and TCD approaches, measurements of relative velocity characteristics in response to different stimuli from rest showed improved but variable agreement, with the strongest correlations observed for the change in average velocity between rest and exercise. Therefore, in addition to absolute measurements, incorporation of relative changes in velocity in response to physiological stimuli is an important consideration for future research using either TCD ultrasound or CFD cerebrovasculature simulations.

## ACKNOWLEDGEMENTS

We acknowledge the resources provided by the Pawsey Supercomputing Centre with funding from the Australian Government and the Government of Western Australia. D.J.G. is supported by a NHMRC Principal Research Fellowship (APP1080914). There are no conflicts of interest.

## Author Contributions

H.T.C., H.J.T., D.J.G., and B.J.D. conceived and designed research; H.J.T., K.J.S. and D.J.G. performed experiments; H.T.C., L.J.K. and B.J.D. developed simulations; H.T.C., H.J.T., L.J.K., K.J.S., D.J.G., and B.J.D. interpreted results of experiments; H.T.C, B.J.D. and D.J.G. drafted manuscript; H.T.C., H.J.T., L.J.K., K.J.S., B.J.D. and D.J.G. edited and revised manuscript; H.T.C., H.J.T., L.J.K., K.J.S., B.J.D. and D.J.G. approved final version of manuscript; H.T.C., L.J.K., and B.J.D. analyzed data; H.T.C., L.J.K., B.J.D. and D.J.G. prepared figures.

