## Supplementary Material for "Comparison of Transcranial Doppler Ultrasound with Computational Fluid Dynamics: Responses to Physiological Stimuli"

HARRISON T. CADDY[1,2]

HANNAH J. THOMAS[3]

LACHLAN J. KELSEY[1,2]

KURT J. SMITH[3,4]

BARRY J. DOYLE[1,2,5,6*]

DANIEL J. GREEN[3*]

[1]*Vascular Engineering Laboratory*, Harry Perkins Institute of Medical Research, Queen Elizabeth II Medical Centre, Nedlands, Australia and the UWA Centre for Medical Research, The University of Western Australia, Perth, Australia
[2]*School of Engineering*, The University of Western Australia, Perth, Australia
[3]*School of Human Sciences (Exercise and Sport Sciences)*, The University of Western Australia, Perth, Australia
[4]*Integrative Physiology Laboratory*, Department of Kinesiology and Nutrition, University of Illinois, Chicago, Illinois
[5]*Australian Research Council Centre for Personalised Therapeutics Technologies*, Melbourne, Australia
[6]*British Heart Foundation Centre for Cardiovascular Science*, The University of Edinburgh, Edinburgh, United Kingdom

* Joint senior authors

**SHORT TITLE**: Comparison of TCD and CFD in response to stimuli

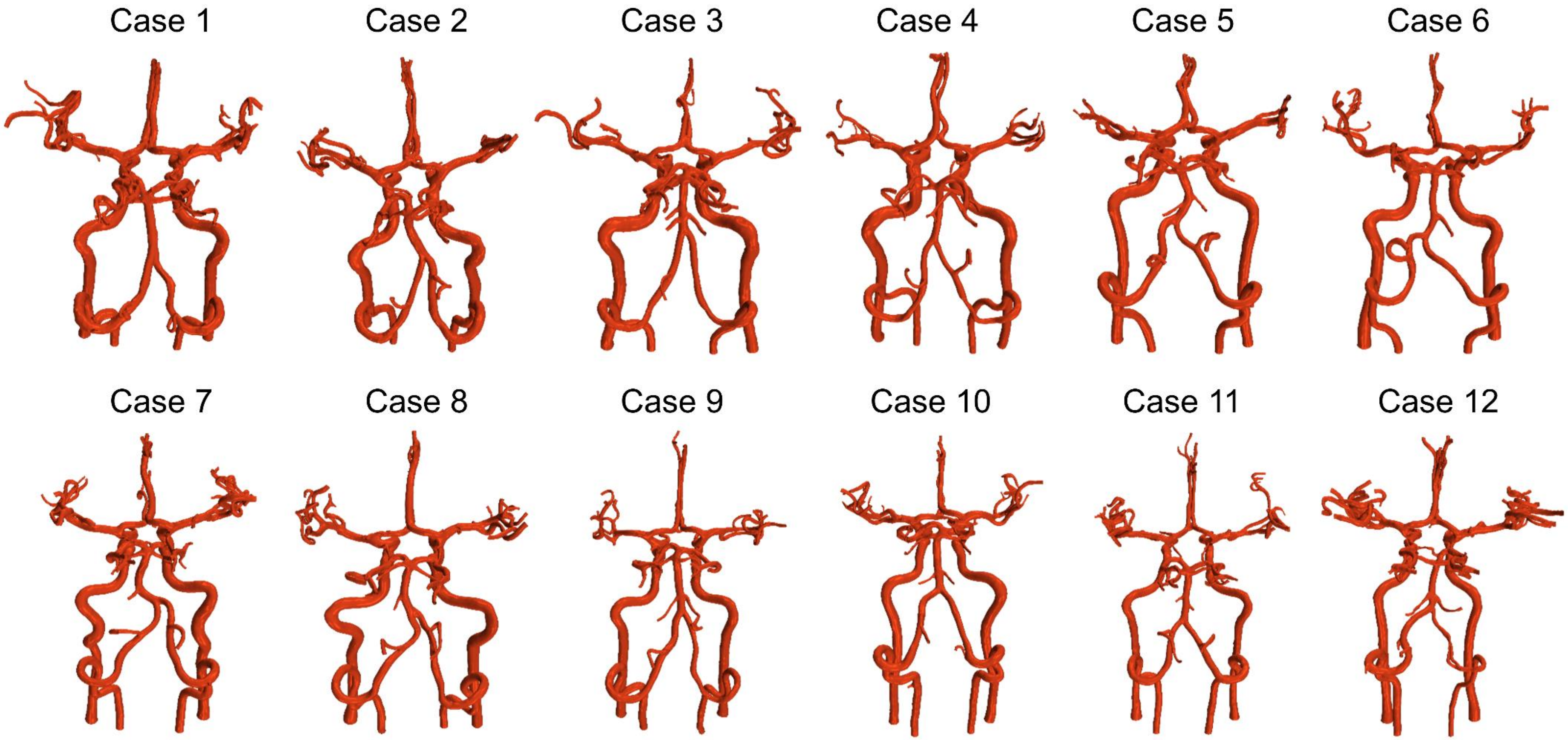


**Figure S1. 3D reconstructions of each case.** All cases exhibited a complete CoW.

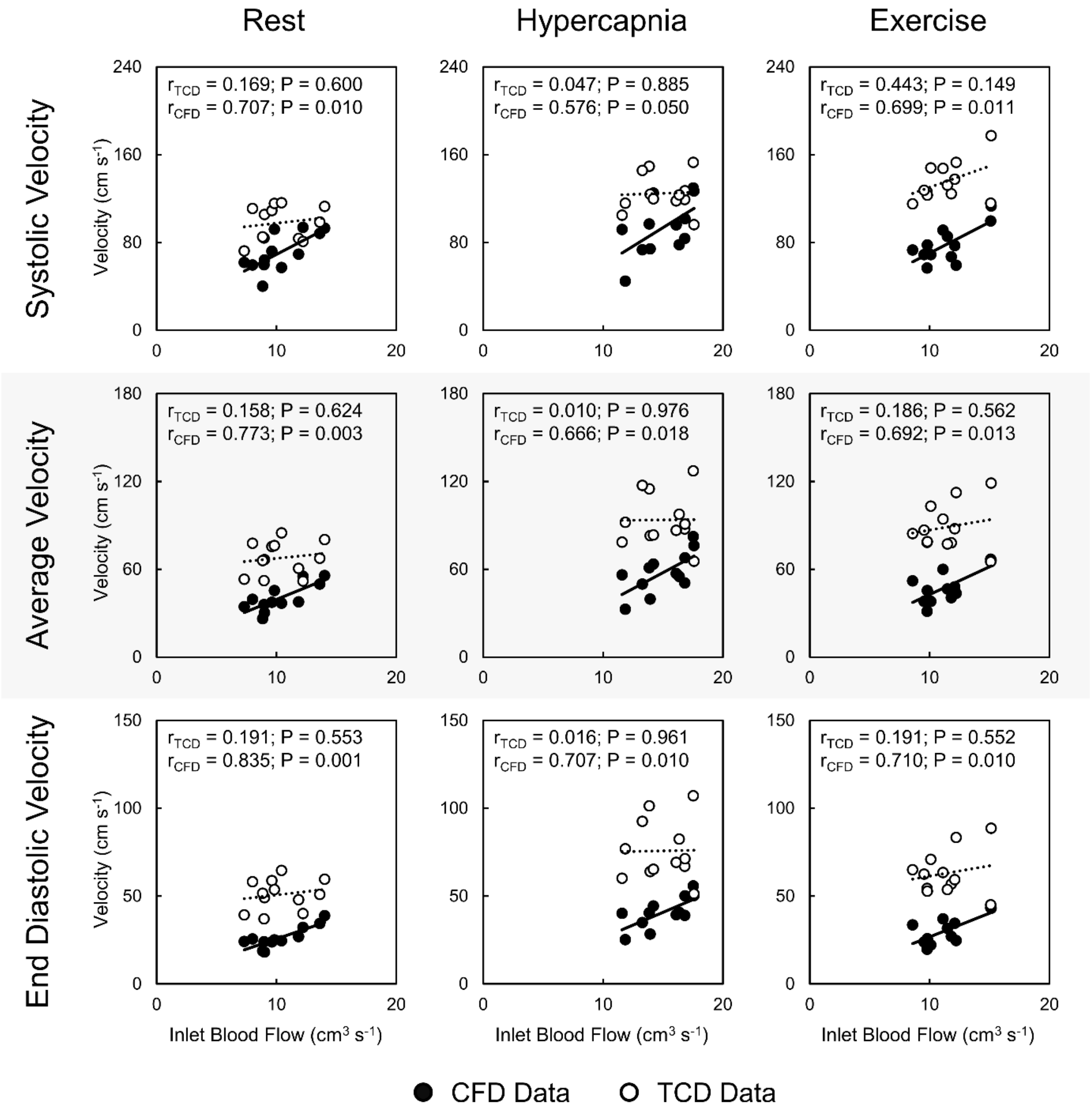


**Figure S2. Correlation plots of total average inlet blood flow with the velocity waveform characteristics of systolic, average and end diastolic maximal velocity extracted from CFD (filled; n=12; 6 male, 6 female) and three cycle averaged TCD (hollow; n=12; 6 male, 6 female) data in the right M1 segment for the conditions of rest, hypercapnia and exercise.**

Total average inlet blood flow was calculated as the sum of the average of the inlet flow waveforms, which were calculated from ultrasound data of velocity and corresponding diameter variation waveforms in the left and right internal carotid and vertebral arteries. Pearson's correlation coefficient (r) and p-value (P) are displayed for each correlation plot.

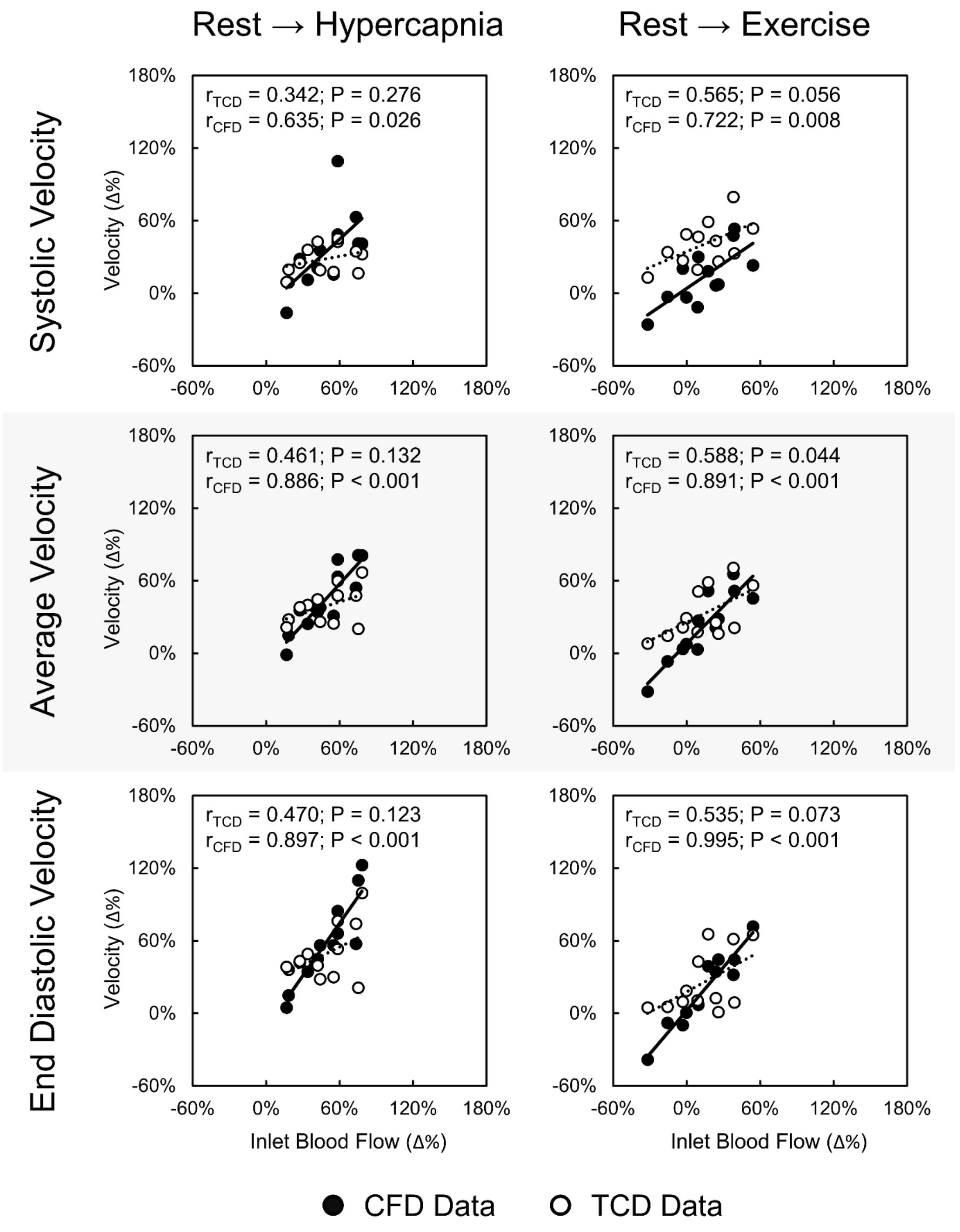


**Figure S3: Correlation plots of the relative change (Δ%) in total average inlet blood flow with the relative change in velocity waveform characteristics (systolic, average and end diastolic velocity) extracted from CFD (filled; n=12; 6 male, 6 female) and three cycle averaged TCD (hollow; n=12; 6 male, 6 female) data in the right M1 segment for responses from rest to hypercapnia (left) and rest to exercise (right).**

Total average inlet blood flow was calculated as the sum of the average of the inlet flow waveforms, which were calculated from ultrasound data of velocity and corresponding diameter variation waveforms in the left and right internal carotid and vertebral arteries. Pearson's correlation coefficient (r) and p-value (P) are displayed for each correlation plot.
